# Somatosensory versus cerebellar contributions to proprioceptive changes associated with motor skill learning: A theta burst stimulation study

**DOI:** 10.1101/2020.08.06.239707

**Authors:** Jasmine L. Mirdamadi, Hannah J. Block

## Abstract

**Background:** It is well established that proprioception (position sense) is important for motor control, yet its role in motor learning and associated plasticity is not well understood. We previously demonstrated that motor skill learning is associated with enhanced proprioception and changes in sensorimotor neurophysiology. However, the neural substrates mediating these effects are unclear.

**Objective:** To determine whether suppressing activity in the cerebellum and somatosensory cortex (S1) affects proprioceptive changes associated with motor skill learning.

**Methods:** 54 healthy young adults practiced a skill involving visually-guided 2D reaching movements through an irregular-shaped track using a robotic manipulandum with their right hand. Proprioception was measured using a passive two-alternative choice task before and after motor practice. Continuous theta burst stimulation (cTBS), was delivered over S1 or the cerebellum (CB) at the end of training for two consecutive days. We compared group differences (S1, CB, Sham) in proprioception and motor skill, quantified by a speed-accuracy function, measured on a third consecutive day (retention).

**Results:** As shown previously, the Sham group demonstrated enhanced proprioceptive sensitivity after training and at retention. The S1 group had impaired proprioceptive function at retention through online changes during practice, whereas the CB group demonstrated offline decrements in proprioceptive function. All groups demonstrated motor skill learning, however, the magnitude of learning differed between the CB and Sham groups, consistent with a role for the cerebellum in motor learning.

**Conclusion:** Overall, these findings suggest that the cerebellum and S1 are important for distinct aspects of proprioceptive changes during skill learning.

## 1. Introduction

Motor learning involves changes in behavior associated with practice. Learning can describe modifications of already-well learned movements (motor adaptation) or the acquisition of new skills (skill learning). Both are associated with changes in motor brain regions, including the motor cortex and cerebellum. These changes can occur during the acquisition phase of learning (i.e., online learning), or between sessions (i.e., offline learning) (Dayan & Cohen, 2011; Kantak & Winstein, 2012).

The extent to which specific brain regions are involved in learning depends on the phase and type of learning. Adaptation involves modification of already well-learned movements to compensate for an external perturbation. This trial-by-trial reduction in errors occurs within minutes (Bastian, 2008; Krakauer & Mazzoni, 2011). Transcranial direct current stimulation (tDCS) research suggests that the cerebellum contributes to the rate of adaptation whereas the primary motor cortex (M1) contributes to retention after the perturbation is removed (Galea et al., 2011).

M1 and the cerebellum likely also contribute to different phases of skill learning (Spampinato & Celnik, 2017). Unlike adaptation, skill learning involves the acquisition of new movement patterns in the absence of a perturbation. There is an improvement in movement quality rather than regaining a baseline performance; this can occur on a longer time scale (i.e., days, weeks, years) (Krakauer & Mazzoni, 2011; Shmuelof et al., 2012). Excitatory cerebellar tDCS facilitated total skill learning through online rather than offline changes (Cantarero et al., 2015). Other studies using the same skill task found that excitatory M1 tDCS enhanced skill retention, without affecting online gains (Reis et al., 2009; Saucedo Marquez et al., 2013). Overall, previous literature suggests a potential dichotomy in the roles of the cerebellum and M1 during adaptation and skill learning, with the cerebellum playing a larger role in online changes and M1 playing a larger role in offline processes.

While the motor brain regions have been investigated frequently in the context of motor learning, sensory functions have received less attention. Given anatomical, functional, and physiological evidence suggesting reciprocal links between sensory and motor processes (Ostry & Gribble, 2016), consolidation of sensory memories may also play a role in motor learning. We and others have shown that motor skill learning is associated with improvements in body position sense (proprioception) that are retained at least 24 hours after practice ends (Cuppone et al., 2018; Mirdamadi & Block, 2020). We also demonstrated changes in somatosensory projections to motor cortex after training (Mirdamadi & Block, 2020). However, the neural substrates driving these sensory changes have not been directly investigated.

Here we consider two possible neural substrates that may be important for proprioceptive changes associated with motor skill learning: the cerebellum and primary somatosensory cortex (S1). Both regions process proprioceptive information from the periphery, and are interconnected with other cortical areas important for sensorimotor integration (Edwards et al., 2019; Gilman, 2002; Ostry & Gribble, 2016; Proske & Gandevia, 2012). Cerebellar involvement in proprioception is thought to be more non-conscious, contributing to online movement corrections (Baumann et al., 2015; Riemann & Lephart, 2002). In contrast, S1 is thought to be involved in higher-order proprioceptive processing and conscious limb detection (Johnson et al., 2008). Here, we applied continuous theta burst stimulation (cTBS), an inhibitory non-invasive brain stimulation paradigm, over S1 or the cerebellum after training. We compared these groups to a Sham group to determine whether S1 and the cerebellum have different roles in proprioceptive changes associated with motor skill learning.

## 2. Methods

### 2.1. Participants

54 right-handed healthy young adults (35 female, age 18-33 years), with no known neurological disorders nor contraindications to transcranial magnetic stimulation (TMS), gave written informed consent and participated. The study was approved by the Indiana University Institutional Review Board.

### 2.2. Experimental Design

Participants were randomly assigned to one of three stimulation groups: S1, Cerebellum (CB), and Sham (Fig.1A). Each group participated in three consecutive sessions that involved motor skill and proprioceptive tasks identical to our previous experiment (Mirdamadi & Block, 2020), with cTBS delivered at the end of day 1 and day On day 3, we assessed retention of proprioceptive function and motor skill in the absence of cTBS. (Fig.1B).

Behavioral tasks were performed using the KINARM Endpoint 2D robotic manipulandum (BKIN). Participants grasped the manipulandum with their right hand and viewed a task display that appeared in the plane of the manipulandum. They performed all tasks without direct vision of their arms or the manipulandum (Mirdamadi & Block, 2020).

For the motor skill, participants navigated a visual cursor representing their hand position (white circle, 10mm diameter) through an irregular shaped track (20×20 cm space, 1.5 cm width) (Fig. 2B) as accurately as possible within the desired movement time range. After each movement, participants received feedback on their speed (too slow, too fast, good speed) and accuracy in the form of points. They were instructed to first prioritize speed, and then improve accuracy (McGrath & Kantak, 2016; Mirdamadi & Block, 2020).

The motor skill assessment on day 1 and day 3 (Fig. 1B) assessed a speed-accuracy tradeoff over five movement time (MT) ranges, with 10 trials in each range, randomized across participants (MT1: 300-600 ms; MT2: 600-850 ms; MT3: 850-1100 ms; MT4: 1100-1400 ms: MT5: 1400-1700 ms). During motor training on day 1 and day 2 (Fig. 1B), participants trained at a fixed MT range (MT3) for 120 trials and 150 trials, respectively (Mirdamadi & Block, 2020).

Proprioception was assessed using a passive two-alternative forced choice task that required subjects to verbally report where their hand was in relation to a constant visual reference marker (Fig. 2D). Proprioception was assessed in the horizontal (left/right) and sagittal (up/down) dimensions, with order randomized across participants. After the hand was moved to the reference, there was a random distractor movement to minimize learning (Mirdamadi & Block, 2020; Wong et al., 2011), followed by a subsequent movement to a test position (Mirdamadi & Block, 2020; Wilson et al., 2010). Test positions followed an adaptive staircase algorithm based on the Parameter Estimation by Sequential Testing method (PEST) (Taylor & Creelman, 1967). There was a total of four staircases, beginning 6 cm left/right or up/down of the reference. Subsequent test positions were adjusted depending upon the subject’s response. Initial step size was 2 cm, and decreased by half when the subject’s response reversed (i.e. from left to right). Each staircase terminated after four reversals (Mirdamadi & Block, 2020).

### 2.3. Transcranial Magnetic Stimulation (TMS) and Recordings

Participants were seated with their arms relaxed on a pillow. TMS was delivered using a Magstim Super Rapid Plus stimulator with a D70^2^ 70-mm figure-of-eight coil (Magstim Company LTD, United Kingdom). BrainSight neuronavigation system (Rogue Research, Montreal, Canada) was used for consistent coil positioning. Surface electromyography (EMG) was recorded from the right first dorsal interosseous (FDI) muscle. EMG signals were amplified (AMT-8; Bortec Biomedical, Calgary, Canada), band-pass filtered (10-1000 Hz), sampled at 5000 Hz, and recorded using Signal software (Cambridge Electronic Design Ltd, United Kingdom).

At the beginning of day 1 and day 2, single pulse TMS was used to identify the FDI “hotspot”, or optimal scalp position that evoked the largest and most consistent motor evoked potential (MEP) in right FDI muscle. The coil was held tangentially with the handle 45° to the midline to evoke posterior-to-anterior current in the cortex. Next, we found resting motor threshold (RMT), defined as the minimum intensity that evoked an MEP at least 50 microvolts in at least 10 out of 20 trials (Rossini et al., 2015).

cTBS was delivered at the end of day 1 and day 2 (Fig. 1B). The S1 target was defined 1 cm posterior and 2 cm lateral to the FDI hotspot (Holmes et al., 2019), with the handle 45° from the midline. The CB target was 3 cm lateral and 1 cm inferior the inion (Casula et al., 2016; Del Olmo et al., 2007; Harrington & Hammond-Tooke, 2015; Koch et al., 2008), with the handle pointing superiorly. cTBS consisted of three pulses presented at 50 Hz, repeated at 5 Hz for 40s, for a total of 600 stimuli (Huang et al., 2005). The intensity was 70% of RMT (Gentner et al., 2008; Goldsworthy et al., 2014). The Sham group experienced cTBS with the coil tilted 90° away from the CB target with only the coil edge on the scalp (Brusa et al., 2012; Monaco et al., 2014).

### 2.4. Data Analysis

For each trial of motor skill (Fig. 2C), we calculated movement time (MT) and percentage of movement trajectory inside the track (in-track accuracy). Only trials of the correct MT were analyzed. For each proprioception assessment, we calculated the proportion of trials that a participant responded left (horizontal dimension) or down (sagittal dimension) across different test positions. Data were fitted with a logistic function upon which bias and sensitivity were calculated. Bias (perceptual boundary) was defined as the 50% point of the fitted function. Since we were interested in detecting improvements in bias independent of direction, we used the absolute bias value in group analyses. Sensitivity (uncertainty) was defined as the distance between the 25% and 75% points of the fitted function (Mirdamadi & Block, 2020; Wilson et al., 2010; Wong et al., 2011) (Fig. 2E).

Since total change in proprioceptive function between baseline and retention (proprioception day 3 – pre day 1) can be manifested through online changes, offline changes, or a combination, we also calculated online and offline changes in bias and sensitivity. Online change was calculated by: (proprioception post day 1 – pre day 1) + (post day 2 – pre day 1). Offline change was calculated by: (proprioception pre day 2 – post day 1) + (day 3 – post day 2).

### 2.5. Statistical Analysis

To determine the effect of cTBS on proprioceptive changes associated with training, we performed a 3-way mixed measures ANOVA with within-subject factors Training Day (day 1, day 2) and Time (pre-training, post-training), and between-subject factor Group (S1, CB, Sham). We ran a one-way ANOVA on each of online changes, offline changes, and total changes. Horizontal and sagittal dimensions were analyzed separately.

To determine the effect of cTBS on motor skill learning, we performed a mixed measures ANOVA with within-subject factors Session (baseline, retention) and MT Bin (MT1, MT2, MT3, MT4, MT5) and between-subject factor Group (S1, CB, Sham) on in-track accuracy.

To assess whether the three groups were similar at baseline, we performed one-way ANOVAs with between-subject factor Group (S1, CB, Sham) on proprioceptive bias and sensitivity. A two-way ANOVA with between-subject factor Group and within-subject factor MT Bin (MT1, MT2, MT3, MT4, MT5) on in-track accuracy was performed to determine baseline skill. Finally, to see if RMT differed across groups or days, we performed a two-way ANOVA with between-subject factor Group and within-subject factor Training Day (day 1, day 2).

For all ANOVAs, assumptions for normality and homogeneity of variance were checked using the Shapiro-Wilk test and Levene’s test, respectively, and log-transformed if necessary. However, all data is plotted using the non-transformed values for clarity. Results were Greenhouse-Geisser corrected if the assumption of sphericity was violated. Significant effects were followed by post-hoc contrasts and corrected for multiple comparisons using Tukey’s HSD method.

## 3. Results

### 3.1. Proprioceptive sensitivity - horizontal dimension

In the horizontal dimension, proprioceptive sensitivity changed differently between timepoints across the three groups, as indicated by a Group x Time interaction [F(2,51) = 4.11, p = 0.022]. No other effects or interactions were significant. The Group x Time interaction reflects a trend for the S1 group having worse, and the CB group better, sensitivity post-training compared to pre-training (S1: t(51) = −1.99, p = 0.053; CB: t(51) = 1.80, p = 0.078). Sham did not show significant changes across time (p > 0.8) (Fig. 3A). After collapsing across Training Day, Sham did not differ significantly from CB or S1, but CB and S1 differed from each other [Group x Time interaction: F(1,34) = 9.76, p = 0.0041], with the S1 group having worse sensitivity after training compared to the CB group.

Total change in horizontal sensitivity, from baseline to retention, differed across groups (F(2,51) = 5.94, p = 0.0054). This reflected worsening for the S1 group compared to Sham (t(51) = 3.44, p = 0.003; Fig. 3C), while the CB group did not differ significantly from Sham or S1 (p > 0.16). Group differences in online proprioceptive changes (F(2,51) = 4.11, p = 0.022) were driven by a difference between the S1 and CB groups (t(51) = 2.85, p = 0.017), with the S1 group getting relatively worse. Sham did not differ significantly from either S1 or CB (p > 0.2). Finally, group differences in offline proprioceptive changes (F(2,51) = 3.68, p = 0.032) were driven by the CB group having worsened offline compared to Sham (t(51) = 2.71, p = 0.024). The S1 group did not differ from the CB or Sham groups (p > 0.3) (Fig. 3B). In summary, the S1 group had worsened total sensitivity in the horizontal dimension relative to Sham, primarily driven by online changes. In contrast, the CB group demonstrated offline decrements compared to Sham.

### 3.2. Proprioceptive sensitivity – sagittal dimension

In the sagittal dimension, sensitivity changed after training similarly across groups, as indicated by a main effect of Time [F(1,51) = 5.33, p = 0.025]. This reflects a lower (better) sensitivity after training compared to before training regardless of Group or Training Day. There was also a trend for a main effect of Training Day [F(1,51) = 3.89, p = 0.054]. This reflects lower (better) sensitivity on day 2 compared to day 1 (Fig. 3C). No other effects or interactions were significant.

Total change in sagittal sensitivity from baseline to retention tended to differ among the three groups (F(2,51) = 3.17, p = 0.0504). Improvement was smallest for the CB group and largest for the Sham group. Neither online or offline changes in sagittal sensitivity differed across groups (p > 0.6) (Fig. 3D).

### 3.3. Proprioceptive bias

Neither horizontal nor sagittal bias was modulated across training days or at retention, nor was it affected by stimulation site. Horizontal bias across training days was not normally distributed and therefore log-transformed. There were no significant interactions or main effects, suggesting that horizontal bias was similar across training days and between groups (all p>0.11, Fig. 4A). At retention, total change in horizontal bias was not different across groups (p>0.5). There were no between-group differences in online or offline changes in horizontal bias (p > 0.2, Fig. 4B).

The three groups were similar across training in sagittal bias, as indicated by the absence of any interactions or main effects (all p>0.1, Fig. 4C). There were no group differences in total sagittal bias change nor in online or offline changes (all p > 0.3, Fig. 4D).

### 3.4. Motor Skill Learning

All groups were able to learn the motor skill, as indicated by a main effect of Session (F(1,51) = 48.55, p < 0.0001). There was a Session x MT Bin interaction [F(4,204)=7.52, p<0.0001, Greenhouse-Geisser corrected], with higher accuracy at retention compared to baseline for all MT bins except MT5 (MT1-MT4: t(240) = −3.92, p = 0.0001; t(240) = −7.78, p<0.0001; t(240) = −3.95, p=0.0001; t(240) = −2.58, p=0.011; MT5: p=0.43) (Fig. 5A). However, a significant Group x Session interaction [F(2,51) = 3.27, p = 0.046] suggests differences in learning across the three groups. The CB group learned significantly less than Sham (t(51) = 2.50, p = 0.041). The Sham group learned the most (6.89% gain, t(51) = −6.02, p = 0.0001) whereas the CB group learned the least (2.85% gain, t(51) = −2.49, p = 0.016). The S1 group gained 4.08% (t(51) = −3.57, p = 0.0008). No other group differences were noted (p>0.2) (Fig. 5B).

### 3.5. Baseline performance and neurophysiology measures

Proprioceptive bias and sensitivity at baseline did not differ between groups in either dimension (all p>0.35). For baseline motor skill, the absence of any effect or interaction involving Group suggests the groups had a similar level of skill before training. A main effect of MT bin [F(4,204) = 282.89, p<0.0001] reflects greater accuracy at slower speeds, as expected. Finally, RMT analysis revealed no significant effects or interactions (all p>0.16).

## 4. Discussion

We compared cerebellar versus S1 contributions to proprioceptive changes associated with motor skill learning. Consistent with previous findings, the Sham group demonstrated motor skill learning and improvements in proprioceptive sensitivity after training, which persisted at retention (Mirdamadi & Block, 2020). cTBS over the cerebellum and S1 impaired proprioceptive sensitivity in the horizontal dimension, though during different phases of the learning process: the cerebellum contributed to offline proprioceptive decrements while S1 contributed to online proprioceptive decrements that persisted at retention.

### 4.1. Cerebellar versus somatosensory involvement in proprioceptive function

Proprioceptive information from the periphery ascends along two routes: the dorsal column-medial-lemniscal pathway, which terminates in S1, and the spinocerebellar pathway, which terminates in the spinocerebellum (Proske & Gandevia, 2012). The cerebellum also has extensive parallel loops with cortical areas for additional sensory inputs and motor commands (Wolpert et al., 1998). Patient studies demonstrate the cerebellum contributes to proprioceptive function (Bhanpuri et al., 2013; Weeks et al., 2017). Individuals with cerebellar damage perform similar to controls in passive proprioceptive tasks, but are impaired in active proprioceptive tasks (Bhanpuri et al., 2013). These findings suggest that the cerebellum may be particularly important for sensory prediction of motor commands. In the horizontal dimension, CB cTBS led to offline decrements in proprioceptive sensitivity. In the sagittal dimension, there was a trend for attenuated sensitivity improvements compared to Sham. Overall, these findings support the role of the cerebellum in proprioception. Given that the cerebellum plays more of a role in proprioception for active movements (Bhanpuri et al., 2013), it would be interesting to test whether other group differences would be observed with an active proprioceptive test.

Neuroimaging studies in patients and non-invasive brain stimulation studies in healthy young adults suggest S1 contributes to proprioceptive function (Ben-Shabat et al., 2015; Findlater et al., 2018; Ingemanson et al., 2019; Kumar et al., 2019; Vidoni et al., 2010). Proprioceptive deficits of the finger post-stroke were best predicted by total sensory system injury (S1, secondary somatosensory cortex, and thalamocortical sensory tract) and functional connectivity between secondary somatosensory cortex and M1 (Ingemanson et al., 2019). Similarly, residual sensory function in chronic stroke was related to functional connectivity in sensorimotor networks, specifically between S1, M1, and supplementary motor area (Vahdat et al., 2019). In neurologically-intact individuals, Kumar et al. observed that S1 cTBS impaired proprioceptive sensitivity immediately after stimulation (Kumar et al., 2019). Our findings suggest S1 has more than transient impact, playing a role in consolidation of proprioceptive changes. Like the CB group, the S1 group seemed to have attenuated improvements in sensitivity in the sagittal dimension. More pronounced was that compared to baseline, sensitivity in the horizontal dimension was worse at retention, primarily due to online decrements. This suggests that the cerebellum and S1 may have different contributions (offline versus online) to proprioceptive changes associated with skill learning. These findings may be analogous to previous research suggesting a dichotomy in cerebellar versus M1 contributions to different phases of motor learning (Galea et al., 2011; Spampinato & Celnik, 2017).

To our knowledge, only one other study has directly compared the effects of S1 and cerebellar stimulation on sensory function. Conte et al. found that S1 but not cerebellar TBS affected somatosensory temporal discrimination (Conte et al., 2012). At first glance, if we simply looked at total changes in proprioceptive function, our results would be consistent with Conte et al.’s findings. However, the current study suggests the cerebellum also contributes to proprioceptive function through offline mechanisms.

Regardless of group or day, we did not detect changes in proprioceptive bias. Several studies have observed changes in bias after motor adaptation (Henriques & Cressman, 2012; Ostry et al., 2010; Vahdat et al., 2011). However, there are mixed reports on whether bias changes after learning without a perturbation (Cuppone et al., 2018; Mirdamadi & Block, 2020; Wong et al., 2011). Further research is needed to elucidate which aspects of learning contribute to changes in proprioceptive bias.

### 4.2. Cerebellar versus somatosensory role in motor skill learning

Cerebellar contributions to motor learning are well documented for adaptation paradigms (Bastian, 2008; Martin et al., 1996; Shadmehr & Mussa-Ivaldi, 1994; Tseng et al., 2007). Recent evidence suggests the cerebellum is also involved in skill learning. Shmuelof et al. observed increases in functional connectivity between the cerebellum and motor cortex that were associated with reduced movement variability after three days of practicing an arc-tracing task (Shmuelof et al., 2014). Further, Spampinanto et al. found changes in cerebellar-motor inhibition early in skill learning on a visuomotor pinch force task, but not later phases of learning (Spampinato & Celnik, 2017). Using the same task, excitatory cerebellar tDCS applied during training enhanced total learning through online improvements in accuracy rather than offline learning (Cantarero et al., 2015). The present study had a similar design to Cantarero et al., except we stimulated at the end of practice to interfere with consolidation. Although the CB group still demonstrated skill learning at retention, individuals learned significantly less than Sham, suggesting that the cerebellum contributes to skill learning. Since we did not measure neurophysiology pre and post cTBS, it is difficult to speculate on the underlying mechanisms. However, other reports of cerebellar TBS have observed changes in cerebellar-motor excitability (Popa et al., 2013), motor cortical inhibition (Harrington & Hammond-Tooke, 2015; Koch et al., 2008), and TMS-evoked activity in M1 and posterior parietal cortex (Casula et al., 2016; Harrington & Hammond-Tooke, 2015). Therefore, disrupting the cerebellum likely influenced skill learning via alterations in interconnected cortical areas through dentato-thalamo connections.

One concern with cerebellar stimulation is the potential for stimulating cervical neck roots that in turn drive changes in performance. Our stimulation intensity of 70% RMT is in alignment with other cerebellar TBS studies using 80% of active motor threshold (Casula et al., 2016; Harrington & Hammond-Tooke, 2015; Koch et al., 2008; Popa et al., 2010, 2013), and lower than studies that used 1 Hz repetitive TMS (Del Olmo et al., 2007; Popa et al., 2010). Further, others have performed control experiments stimulating the nerve roots directly, and found cerebellar but not cervical neck muscle stimulation affected cerebellar-motor connectivity and behavior on a tapping task [28,53]. Thus, it is unlikely that our results are due to neck muscle stimulation.

Although most literature has focused on motor contributions to learning, reciprocal connections between motor and somatosensory cortices provide a framework for motor learning to also involve somatosensory plasticity (Ostry & Gribble, 2016). At the behavioral level, proprioception changes after motor adaptation and visuomotor skill learning (Henriques & Cressman, 2012; Mirdamadi & Block, 2020; Ostry et al., 2010; Wong et al., 2011). There is also evidence from somatosensory evoked potentials and resting state functional connectivity to suggest plasticity involving sensory areas following motor adaptation (Nasir et al., 2013; Ostry & Gribble, 2016; Vahdat et al., 2011). Further, inhibitory repetitive TMS over S1 prior to training on a visuomotor tracking task impaired the magnitude of learning compared to Sham (Vidoni et al., 2010). In the present study, the S1 group learned less than Sham, but the difference was not significant. An important distinction between the two studies is the time at which stimulation was delivered. Stimulation before training may have affected proprioception, motor control, or both. In contrast, since we delivered stimulation after training, we rule out any stimulation-induced differences in training performance that may have otherwise affected learning (Kumar et al., 2019).

Kumar et al. (2019) demonstrated that S1 cTBS delivered after force-field learning with a gradual load onset blocked motor memory consolidation (Kumar et al., 2019). At first, our findings seem discrepant with Kumar et.’s findings, but it is difficult to make direct comparisons given the different learning paradigms. Adaptation involves overcoming a perturbation to an already well-learned behavior, with learning indicated by a reduction in systematic errors. In contrast, skill learning involves acquiring new movement patterns without a perturbation, with learning indicated by a shift in the speed-accuracy tradeoff (Krakauer & Mazzoni, 2011; Shmuelof et al., 2012). However, it is important to consider that when learning a new skill, elements of adaptation (i.e. learning the dynamics of a tennis racket) and their associated neural mechanisms may contribute. For instance, cerebellar-motor networks changed during both the early phases of skill learning (i.e. the first training day), as well as following adaptation (Galea et al., 2011; Spampinato & Celnik, 2017, 2018). It is unclear why S1 cTBS abolished retention after adaptation with a gradual load onset but not skill learning as in the present study. One possibility is that skill learning involves a more distributed network compared to adaptation. This hypothesis is consistent with Kumar’s findings that demonstrated partial retention when S1 was suppressed after adaptation with an abrupt load onset. The authors suggested that with an abrupt load, explicit strategies likely require areas other than S1 for learning. Similarly, with skill learning, explicit processes may be involved when subjects explore different strategies to find the optimal movement patterns for completing the maze.

Unfortunately, since the speed-accuracy function was only probed at baseline and retention, we cannot infer anything about online versus offline learning. We did not analyze training performance for two reasons; first, there is a learning-performance distinction such that training performance does not necessarily indicate learning (Kantak & Winstein, 2012). More importantly, changes in performance at a fixed speed may be misrepresentative of total learning which is operationalized by a speed-accuracy tradeoff. If performance at a single speed plateaus during training, it says nothing about how performance changes across the entire speed-accuracy function (Wickelgren, 1977). Future studies will be needed to probe the speed-accuracy function within and between training to assess online versus offline skill learning. Given that the cerebellum and S1 had different online and offline contributions to proprioception, it would be interesting to see whether a similar dichotomy would be observed for skill learning.

## 5. Conclusions

Proprioceptive changes associated with motor skill learning are mediated by both the cerebellum and S1. However, these regions appear to contribute to temporally distinct processes, with cerebellum linked to offline and S1 to online proprioceptive changes. Future research is needed to test whether the cerebellum and somatosensory cortex contribute differently to online versus offline motor skill learning.

**Figure 1A.**
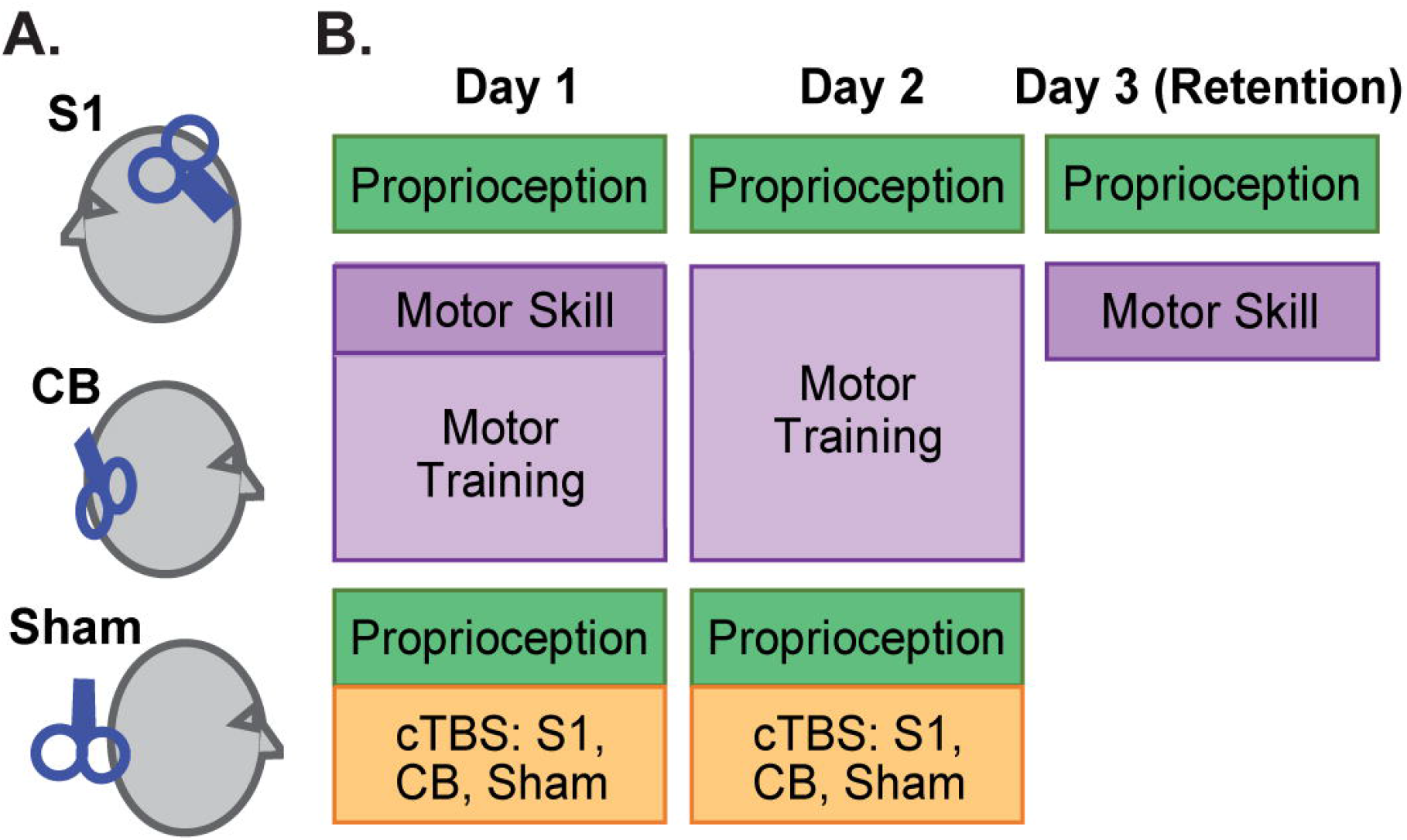
Depiction of stimulation targets used for 3 different groups. Continuous theta burst stimulation (cTBS) was delivered over either the left primary somatosensory cortex (S1), right lateral cerebellum (CB), or right lateral cerebellum with coil tilted away (Sham) **B.** Experimental Design. Proprioceptive function was measured before and after motor training on day 1 and day 2, as in our previous experiment *(Mirdamadi & Block, 2020)*. cTBS was delivered after the behavioral tasks on day 1 and day 2. Retention of proprioceptive function and motor learning was evaluated on day 3 in the absence of cTBS. The motor skill was assessed at 5 different speeds on day 1 and day 3 to evaluate a speed-accuracy trade-off. On day 1 and day 2, motor training was performed at a fixed speed.

**Figure 2A.**
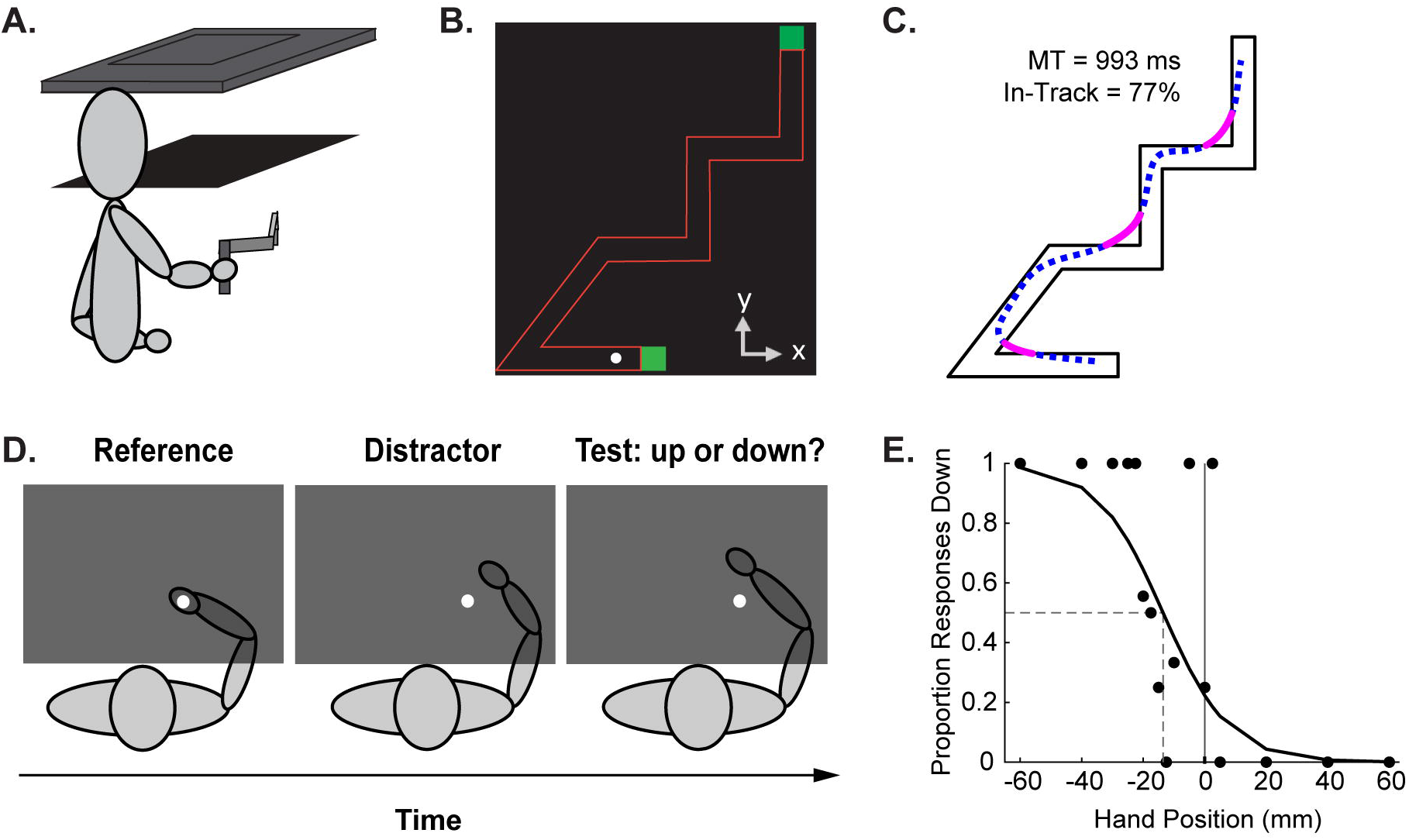
Depiction of 2-D virtual reality apparatus used for the proprioception and motor tasks. Subjects performed the motor skill with their right hand, grasping a robotic manipulandum, and had no vision of their arms. **B.** Bird’s eye view of motor skill task display. Subject was seated in the direction of the negative y-axis, centered with the track. Subjects navigated the white cursor with the robotic manipulandum through the irregular shaped track, moving from the lower green starting square to the upper green end square. **C.** Representative movement trajectory used to compute movement time (MT) and in-track accuracy. Blue dashed line represents parts of the movement path that were inside the track. Magenta line represents parts of the movement path that were outside the track. **D.** Bird’s eye view of passive proprioception assessment. Participants verbally reported the position of their unseen right hand in relation to a visual reference (white circle), located at the center of the motor skill track. Proprioception was assessed in the horizontal dimension, where participants indicated whether their hand was to the left or right of the reference, and sagittal dimension, where participants indicated whether their hand was up or down from the reference. **E.** Example subject proprioceptive data fitted with logistic function. Bias was defined as the 50% point of the fitted function. Sensitivity was defined as the difference between the 25% and 75% points of the fitted function. For this subject, the bias, or perceptual boundary, was computed as −13.55 mm and the sensitivity was 23.83 cm.

**Figure 3.**
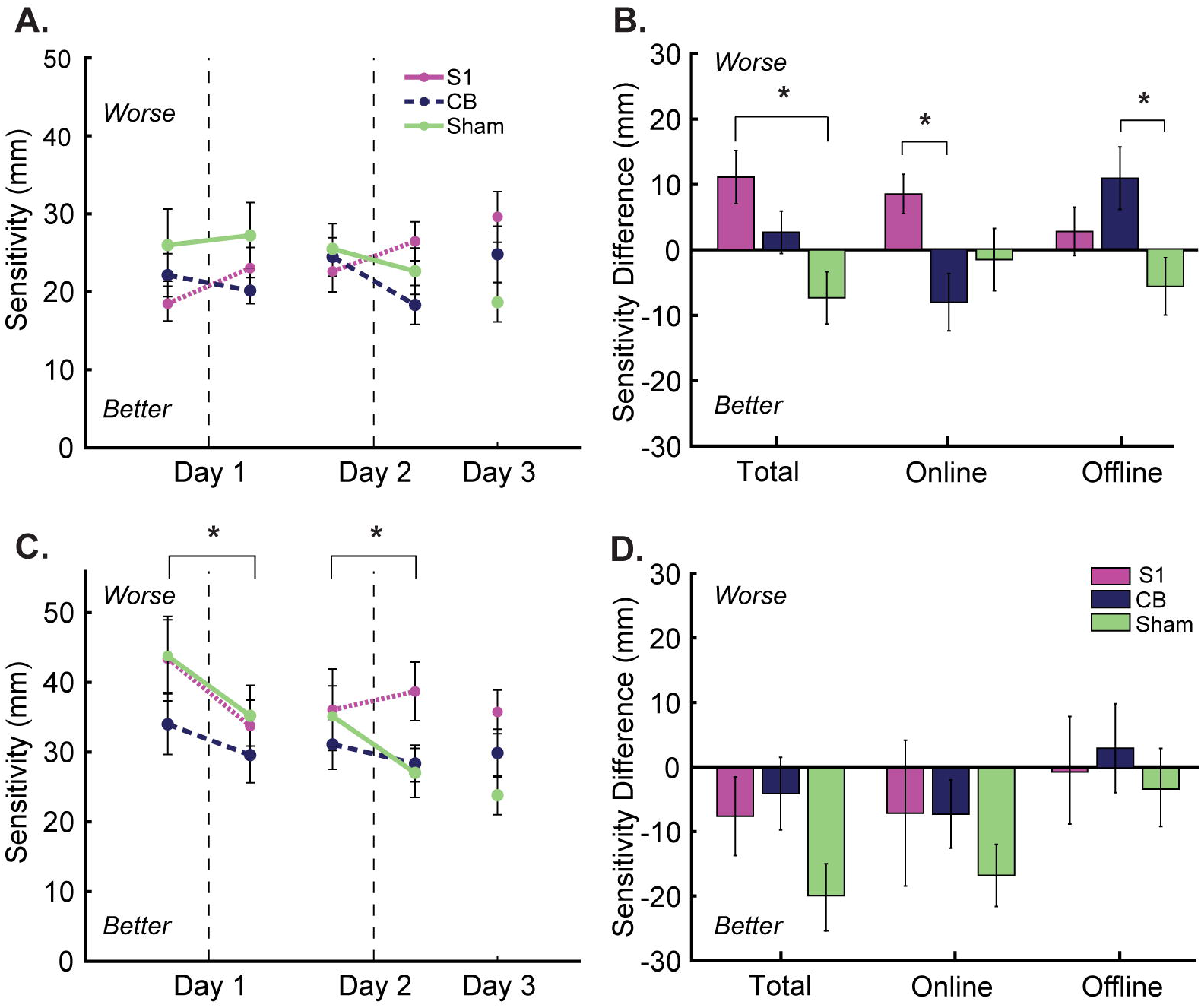
Proprioceptive sensitivity changes in the horizontal dimension (top row) and sagittal dimension (bottom row). Lower values represent better proprioceptive function. **A.** Vertical dashed lines delineate pre-training and post-training on Day 1 and Day 2. Error bars are standard error of the mean. Proprioceptive sensitivity in the horizontal dimension changed differently across time for the S1 and CB groups**. B.** Total changes in proprioceptive function (retention – baseline) broken down into Online and Offline changes. * Tukey pairwise comparison p < 0.05. **C.** Proprioceptive sensitivity in the sagittal dimension decreased (improved) after training regardless of Group or Training Day * denotes main effect of Time, p <0.05. **D.** The magnitude of improvement in total proprioceptive sensitivity was less for the CB and S1 groups compared to Sham, though was not statistically significant (F(2,51) = 3.17, p = 0.0504). There were no group differences in online proprioceptive change or offline proprioceptive change.

**Figure 1.**
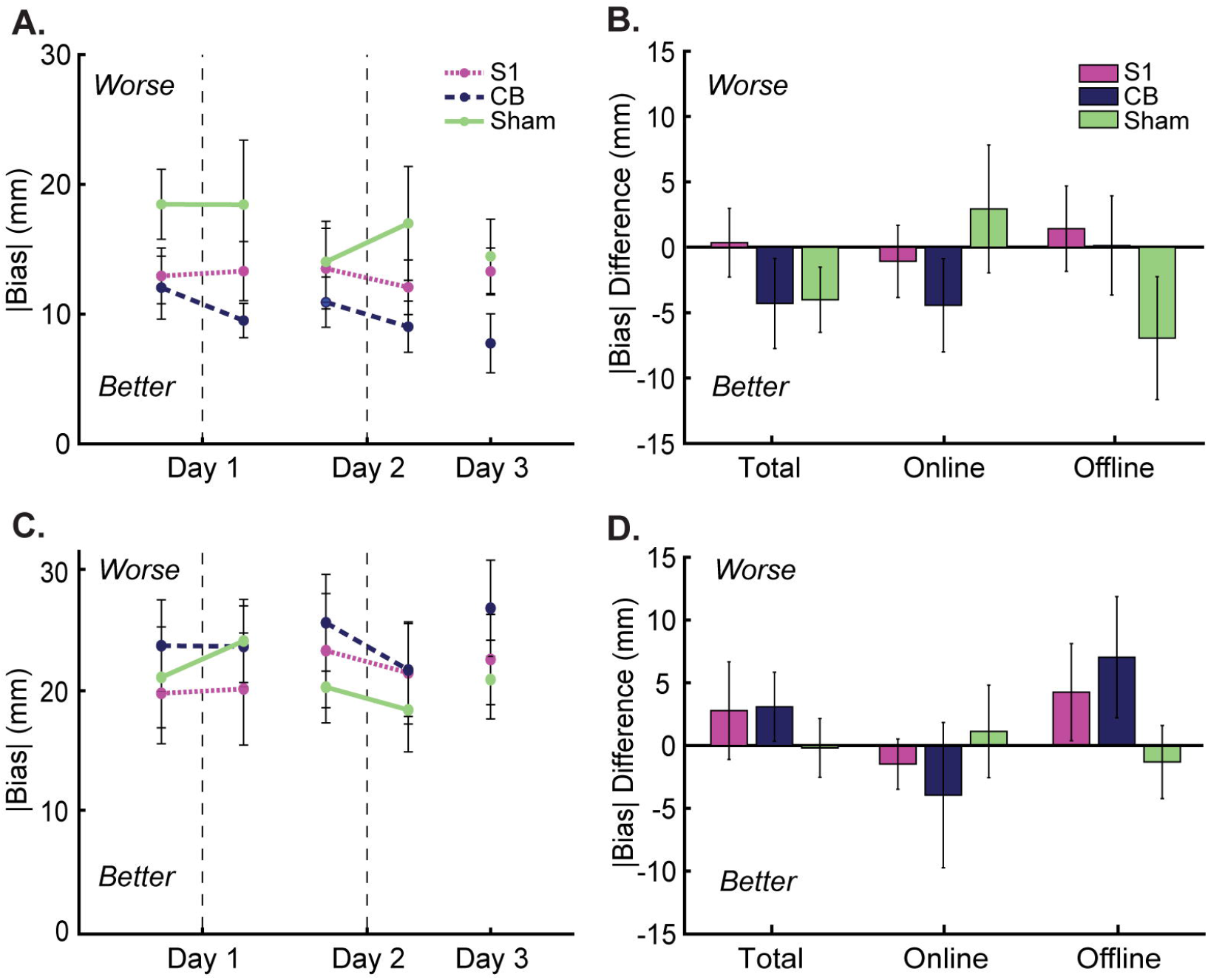
Proprioceptive bias changes in the horizontal dimension (top row) and sagittal dimension (bottom row). Lower values represent better proprioceptive function. Vertical dashed lines delineate pre-training and post-training on Day 1 and Day 2. Error bars are standard error of the mean. **A.** Proprioceptive bias in the horizontal dimension was not modulated differently across groups. **B.** Total, online, and offline changes did not differ across groups. **C.** Proprioceptive bias in the sagittal dimension was not modulated differently across groups. **D.** Total, online, and offline changes did not differ across groups.

**Figure 2A.**
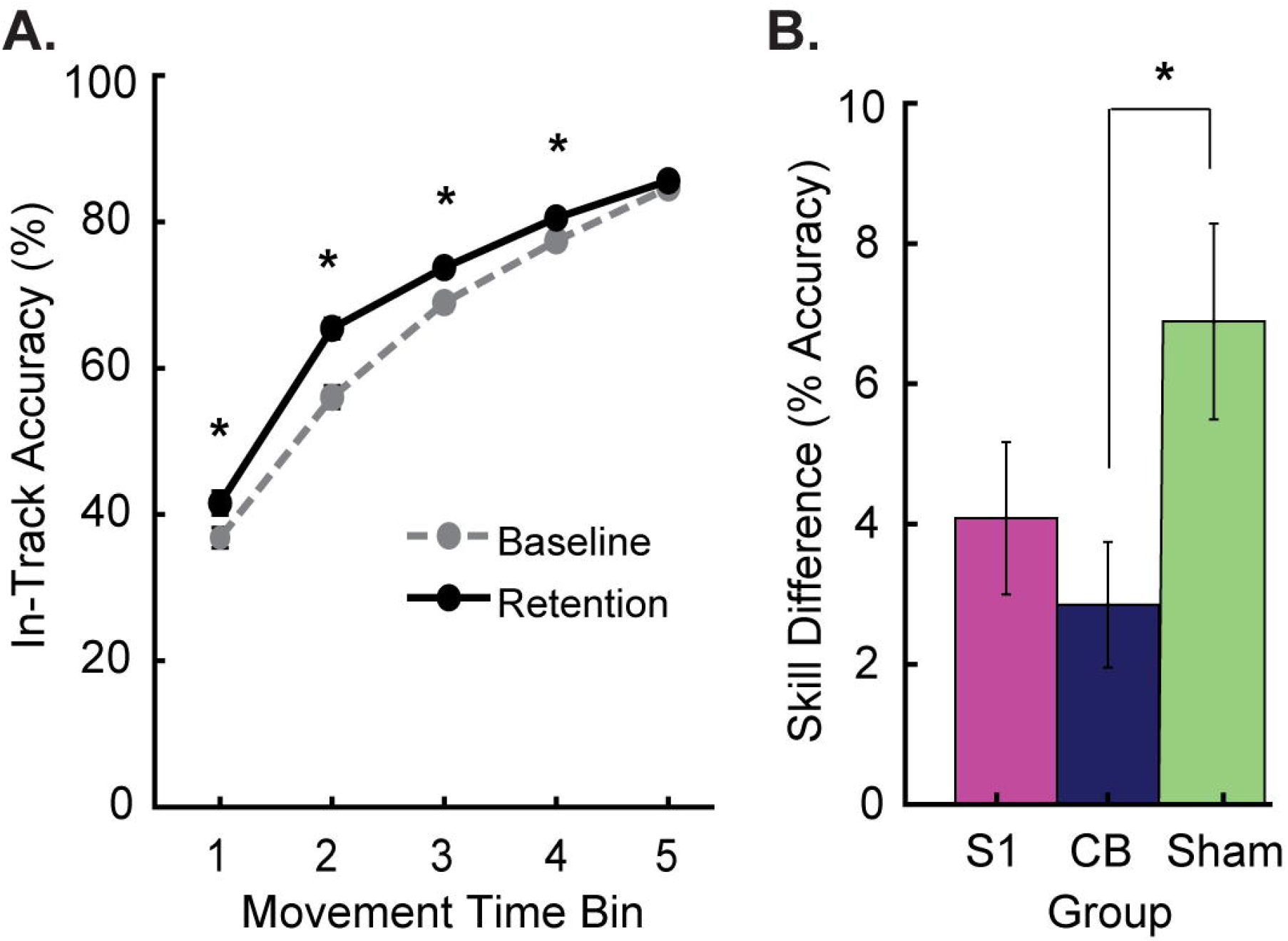
Speed-accuracy function for the motor skill, collapsed across all groups. Higher numbers indicate better performance. The three groups showed evidence of learning at the first four movement time (MT) bins. *p<0.005. **B.** Total skill learning at retention relative to baseline, collapsed across 5 MT bins. The CB group learned to a lesser extent than Sham. *p<0.05.

## Acknowledgements

This work was supported by the National Science Foundation (1342962, JLM; 1753915, HJB).

## References

Bastian, A. J. (2008). Understanding sensorimotor adaptation and learning for rehabilitation: Current Opinion in Neurology, 21(6), 628–633. https://doi.org/10.1097/WCO.0b013e328315a293

Baumann, O., Borra, R. J., Bower, J. M., Cullen, K. E., Habas, C., Ivry, R. B., Leggio, M., Mattingley, J. B., Molinari, M., Moulton, E. A., Paulin, M. G., Pavlova, M. A., Schmahmann, J. D., & Sokolov, A. A. (2015). Consensus Paper: The Role of the Cerebellum in Perceptual Processes. The Cerebellum, 14(2), 197–220. https://doi.org/10.1007/s12311-014-0627-7

Ben-Shabat, E., Matyas, T. A., Pell, G. S., Brodtmann, A., & Carey, L. M. (2015). The Right Supramarginal Gyrus Is Important for Proprioception in Healthy and Stroke-Affected Participants: A Functional MRI Study. Frontiers in Neurology, 6. https://doi.org/10.3389/fneur.2015.00248

Bhanpuri, N. H., Okamura, A. M., & Bastian, A. J. (2013). Predictive Modeling by the Cerebellum Improves Proprioception. Journal of Neuroscience, 33(36), 14301–14306. https://doi.org/10.1523/JNEUROSCI.0784-13.2013

Brusa, L., Ceravolo, R., Kiferle, L., Monteleone, F., Iani, C., Schillaci, O., Stanzione, P., & Koch, G. (2012). Metabolic changes induced by theta burst stimulation of the cerebellum in dyskinetic Parkinson’s disease patients. Parkinsonism & Related Disorders, 18(1), 59–62. https://doi.org/10.1016/j.parkreldis.2011.08.019

Cantarero, G., Spampinato, D., Reis, J., Ajagbe, L., Thompson, T., Kulkarni, K., & Celnik, P. (2015). Cerebellar Direct Current Stimulation Enhances On-Line Motor Skill Acquisition through an Effect on Accuracy. Journal of Neuroscience, 35(7), 3285–3290. https://doi.org/10.1523/JNEUROSCI.2885-14.2015

Casula, E. P., Pellicciari, M. C., Ponzo, V., Stampanoni Bassi, M., Veniero, D., Caltagirone, C., & Koch, G. (2016). Cerebellar theta burst stimulation modulates the neural activity of interconnected parietal and motor areas. Scientific Reports, 6(1). https://doi.org/10.1038/srep36191

Conte, A., Rocchi, L., Nardella, A., Dispenza, S., Scontrini, A., Khan, N., & Berardelli, A. (2012). Theta-Burst Stimulation-Induced Plasticity over Primary Somatosensory Cortex Changes Somatosensory Temporal Discrimination in Healthy Humans. PLoS ONE, 7(3). https://doi.org/10.1371/journal.pone.0032979

Cuppone, A. V., Semprini, M., & Konczak, J. (2018). Consolidation of human somatosensory memory during motor learning. Behavioural Brain Research, 347, 184–192. https://doi.org/10.1016/j.bbr.2018.03.013

Dayan, E., & Cohen, L. G. (2011). Neuroplasticity Subserving Motor Skill Learning. Neuron, 72(3), 443–454. https://doi.org/10.1016/j.neuron.2011.10.008

Del Olmo, M. F., Cheeran, B., Koch, G., & Rothwell, J. C. (2007). Role of the Cerebellum in Externally Paced Rhythmic Finger Movements. Journal of Neurophysiology, 98(1), 145–152. https://doi.org/10.1152/jn.01088.2006

Edwards, L. L., King, E. M., Buetefisch, C. M., & Borich, M. R. (2019). Putting the “Sensory” Into Sensorimotor Control: The Role of Sensorimotor Integration in Goal-Directed Hand Movements After Stroke. Frontiers in Integrative Neuroscience, 13, 16. https://doi.org/10.3389/fnint.2019.00016

Findlater, S. E., Hawe, R. L., Semrau, J. A., Kenzie, J. M., Yu, A. Y., Scott, S. H., & Dukelow, S. P. (2018). Lesion locations associated with persistent proprioceptive impairment in the upper limbs after stroke. NeuroImage: Clinical, 20, 955–971. https://doi.org/10.1016/j.nicl.2018.10.003

Galea, J. M., Vazquez, A., Pasricha, N., Orban de Xivry, J.-J., & Celnik, P. (2011). Dissociating the Roles of the Cerebellum and Motor Cortex during Adaptive Learning: The Motor Cortex Retains What the Cerebellum Learns. Cerebral Cortex, 21(8), 1761–1770. https://doi.org/10.1093/cercor/bhq246

Gentner, R., Wankerl, K., Reinsberger, C., Zeller, D., & Classen, J. (2008). Depression of Human Corticospinal Excitability Induced by Magnetic Theta-burst Stimulation: Evidence of Rapid Polarity-Reversing Metaplasticity. Cerebral Cortex, 18(9), 2046–2053. https://doi.org/10.1093/cercor/bhm239

Gilman, S. (2002). Joint position sense and vibration sense: Anatomical organisation and assessment. Journal of Neurology, Neurosurgery & Psychiatry, 73(5), 473–477. https://doi.org/10.1136/jnnp.73.5.473

Goldsworthy, M. R., Müller-Dahlhaus, F., Ridding, M. C., & Ziemann, U. (2014). Inter-subject Variability of LTD-like Plasticity in Human Motor Cortex: A Matter of Preceding Motor Activation. Brain Stimulation, 7(6), 864–870. https://doi.org/10.1016/j.brs.2014.08.004

Harrington, A., & Hammond-Tooke, G. D. (2015). Theta Burst Stimulation of the Cerebellum Modifies the TMS-Evoked N100 Potential, a Marker of GABA Inhibition. PLOS ONE, 10(11), e0141284. https://doi.org/10.1371/journal.pone.0141284

Henriques, D. Y. P., & Cressman, E. K. (2012). Visuomotor Adaptation and Proprioceptive Recalibration. Journal of Motor Behavior, 44(6), 435–444. https://doi.org/10.1080/00222895.2012.659232

Holmes, N. P., Tamè, L., Beeching, P., Medford, M., Rakova, M., Stuart, A., & Zeni, S. (2019). Locating primary somatosensory cortex in human brain stimulation studies: Experimental evidence. Journal of Neurophysiology, 121(1), 336–344. https://doi.org/10.1152/jn.00641.2018

Huang, Y.-Z., Edwards, M. J., Rounis, E., Bhatia, K. P., & Rothwell, J. C. (2005). Theta burst stimulation of the human motor cortex. Neuron, 45(2), 201–206. https://doi.org/10.1016/j.neuron.2004.12.033

Ingemanson, M. L., Rowe, J. R., Chan, V., Riley, J., Wolbrecht, E. T., Reinkensmeyer, D. J., & Cramer, S. C. (2019). Neural Correlates of Passive Position Finger Sense After Stroke. Neurorehabilitation and Neural Repair, 33(9), 740–750. https://doi.org/10.1177/1545968319862556

Johnson, E. O., Babis, G. C., Soultanis, K. C., & Soucacos, P. N. (2008). Functional neuroanatomy of proprioception. Journal of Surgical Orthopaedic Advances, 17(3), 159–164.

Kantak, S. S., & Winstein, C. J. (2012). Learning–performance distinction and memory processes for motor skills: A focused review and perspective. Behavioural Brain Research, 228(1), 219–231. https://doi.org/10.1016/j.bbr.2011.11.028

Koch, G., Mori, F., Marconi, B., Codecà, C., Pecchioli, C., Salerno, S., Torriero, S., Lo Gerfo, E., Mir, P., Oliveri, M., & Caltagirone, C. (2008). Changes in intracortical circuits of the human motor cortex following theta burst stimulation of the lateral cerebellum. Clinical Neurophysiology, 119(11), 2559–2569. https://doi.org/10.1016/j.clinph.2008.08.008

Krakauer, J. W., & Mazzoni, P. (2011). Human sensorimotor learning: Adaptation, skill, and beyond. Current Opinion in Neurobiology, 21(4), 636–644. https://doi.org/10.1016/j.conb.2011.06.012

Kumar, N., Manning, T. F., & Ostry, D. J. (2019). Somatosensory cortex participates in the consolidation of human motor memory. PLoS Biology, 17(10), e3000469. https://doi.org/10.1371/journal.pbio.3000469

Martin, T. A., Keating, J. G., Goodkin, H. P., Bastian, A. J., & Thach, W. T. (1996). Throwing while looking through prisms: I. Focal olivocerebellar lesions impair adaptation. Brain, 119(4), 1183–1198. https://doi.org/10.1093/brain/119.4.1183

McGrath, R. L., & Kantak, S. S. (2016). Reduced asymmetry in motor skill learning in left-handed compared to right-handed individuals. Human Movement Science, 45, 130–141. https://doi.org/10.1016/j.humov.2015.11.012

Mirdamadi, J. L., & Block, H. J. (2020). Somatosensory changes associated with motor skill learning. Journal of Neurophysiology. https://doi.org/10.1152/jn.00497.2019

Monaco, J., Casellato, C., Koch, G., & D’Angelo, E. (2014). Cerebellar theta burst stimulation dissociates memory components in eyeblink classical conditioning. European Journal of Neuroscience, 40(9), 3363–3370. https://doi.org/10.1111/ejn.12700

Nasir, S. M., Darainy, M., & Ostry, D. J. (2013). Sensorimotor adaptation changes the neural coding of somatosensory stimuli. Journal of Neurophysiology, 109(8), 2077–2085. https://doi.org/10.1152/jn.00719.2012

Ostry, D. J., Darainy, M., Mattar, A. A. G., Wong, J., & Gribble, P. L. (2010). Somatosensory Plasticity and Motor Learning. Journal of Neuroscience, 30(15), 5384–5393. https://doi.org/10.1523/JNEUROSCI.4571-09.2010

Ostry, D. J., & Gribble, P. L. (2016). Sensory Plasticity in Human Motor Learning. Trends in Neurosciences, 39(2), 114–123. https://doi.org/10.1016/j.tins.2015.12.006

Popa, T., Russo, M., & Meunier, S. (2010). Long-lasting inhibition of cerebellar output. Brain Stimulation: Basic, Translational, and Clinical Research in Neuromodulation, 3(3), 161–169. https://doi.org/10.1016/j.brs.2009.10.001

Popa, T., Velayudhan, B., Hubsch, C., Pradeep, S., Roze, E., Vidailhet, M., Meunier, S., & Kishore, A. (2013). Cerebellar Processing of Sensory Inputs Primes Motor Cortex Plasticity. Cerebral Cortex, 23(2), 305–314. https://doi.org/10.1093/cercor/bhs016

Proske, U., & Gandevia, S. C. (2012). The Proprioceptive Senses: Their Roles in Signaling Body Shape, Body Position and Movement, and Muscle Force. Physiological Reviews, 92(4), 1651–1697. https://doi.org/10.1152/physrev.00048.2011

Reis, J., Schambra, H. M., Cohen, L. G., Buch, E. R., Fritsch, B., Zarahn, E., Celnik, P. A., & Krakauer, J. W. (2009). Noninvasive cortical stimulation enhances motor skill acquisition over multiple days through an effect on consolidation. Proceedings of the National Academy of Sciences, 106(5), 1590–1595. https://doi.org/10.1073/pnas.0805413106

Riemann, B. L., & Lephart, S. M. (2002). The Sensorimotor System, Part I: The Physiologic Basis of Functional Joint Stability. Journal of Athletic Training, 37(1), 71–79.

Rossini, P. M., Burke, D., Chen, R., Cohen, L. G., Daskalakis, Z., Di Iorio, R., Di Lazzaro, V., Ferreri, F., Fitzgerald, P. B., George, M. S., Hallett, M., Lefaucheur, J. P., Langguth, B., Matsumoto, H., Miniussi, C., Nitsche, M. A., Pascual-Leone, A., Paulus, W., Rossi, S., … Ziemann, U. (2015). Non-invasive electrical and magnetic stimulation of the brain, spinal cord, roots and peripheral nerves: Basic principles and procedures for routine clinical and research application. An updated report from an I.F.C.N. Committee. Clinical Neurophysiology, 126(6), 1071–1107. https://doi.org/10.1016/j.clinph.2015.02.001

Saucedo Marquez, C. M., Zhang, X., Swinnen, S. P., Meesen, R., & Wenderoth, N. (2013). Task-Specific Effect of Transcranial Direct Current Stimulation on Motor Learning. Frontiers in Human Neuroscience, 7. https://doi.org/10.3389/fnhum.2013.00333

Shadmehr, R., & Mussa-Ivaldi, F. (1994). Adaptive representation of dynamics during learning of a motor task. The Journal of Neuroscience, 14(5), 3208–3224. https://doi.org/10.1523/JNEUROSCI.14-05-03208.1994

Shmuelof, L., Krakauer, J. W., & Mazzoni, P. (2012). How is a motor skill learned? Change and invariance at the levels of task success and trajectory control. Journal of Neurophysiology, 108(2), 578–594. https://doi.org/10.1152/jn.00856.2011

Shmuelof, L., Yang, J., Caffo, B., Mazzoni, P., & Krakauer, J. W. (2014). The neural correlates of learned motor acuity. Journal of Neurophysiology, 112(4), 971–980. https://doi.org/10.1152/jn.00897.2013

Spampinato, D., & Celnik, P. (2017). Temporal dynamics of cerebellar and motor cortex physiological processes during motor skill learning. Scientific Reports, 7, 40715. https://doi.org/10.1038/srep40715

Spampinato, D., & Celnik, P. (2018). Deconstructing skill learning and its physiological mechanisms. Cortex. https://doi.org/10.1016/j.cortex.2018.03.017

Taylor, M. M., & Creelman, C. D. (1967). PEST: Efficient Estimates on Probability Functions. The Journal of the Acoustical Society of America, 41(4A), 782–787. https://doi.org/10.1121/1.1910407

Tseng, Y., Diedrichsen, J., Krakauer, J. W., Shadmehr, R., & Bastian, A. J. (2007). Sensory Prediction Errors Drive Cerebellum-Dependent Adaptation of Reaching. Journal of Neurophysiology, 98(1), 54–62. https://doi.org/10.1152/jn.00266.2007

Vahdat, S., Darainy, M., Milner, T. E., & Ostry, D. J. (2011). Functionally specific changes in resting-state sensorimotor networks after motor learning. The Journal of Neuroscience: The Official Journal of the Society for Neuroscience, 31(47), 16907–16915. https://doi.org/10.1523/JNEUROSCI.2737-11.2011

Vahdat, S., Darainy, M., Thiel, A., & Ostry, D. J. (2019). A Single Session of Robot-Controlled Proprioceptive Training Modulates Functional Connectivity of Sensory Motor Networks and Improves Reaching Accuracy in Chronic Stroke. Neurorehabilitation and Neural Repair, 33(1), 70–81. https://doi.org/10.1177/1545968318818902

Vidoni, E. D., Acerra, N. E., Dao, E., Meehan, S. K., & Boyd, L. A. (2010). Role of the primary somatosensory cortex in motor learning: An rTMS study. Neurobiology of Learning and Memory, 93(4), 532–539. https://doi.org/10.1016/j.nlm.2010.01.011

Weeks, H. M., Therrien, A. S., & Bastian, A. J. (2017). Proprioceptive Localization Deficits in People With Cerebellar Damage. The Cerebellum, 16(2), 427–437. https://doi.org/10.1007/s12311-016-0819-4

Wickelgren, W. A. (1977). Speed-accuracy tradeoff and information processing dynamics. Acta Psychologica, 41(1), 67–85. https://doi.org/10.1016/0001-6918(77)90012-9

Wilson, E. T., Wong, J., & Gribble, P. L. (2010). Mapping Proprioception across a 2D Horizontal Workspace. PLoS ONE, 5(7), e11851. https://doi.org/10.1371/journal.pone.0011851

Wolpert, D. M., Miall, R. C., & Kawato, M. (1998). Internal models in the cerebellum. Trends in Cognitive Sciences, 2(9), 338–347. https://doi.org/10.1016/S1364-6613(98)01221-2

Wong, J. D., Wilson, E. T., & Gribble, P. L. (2011). Spatially selective enhancement of proprioceptive acuity following motor learning. Journal of Neurophysiology, 105(5), 2512–2521. https://doi.org/10.1152/jn.00949.2010

